# Self-supervised learning to predict intrahepatic cholangiocarcinoma transcriptomic classes on routine histology

**DOI:** 10.1101/2024.01.15.575652

**Authors:** Aurélie Beaufrère, Tristan Lazard, Rémy Nicolle, Gwladys Lubuela, Jérémy Augustin, Miguel Albuquerque, Baptiste Pichon, Camille Pignolet, Victoria Priori, Nathalie Théou-Anton, Mickael Lesurtel, Mohamed Bouattour, Kévin Mondet, Jérôme Cros, Julien Calderaro, Thomas Walter, Valérie Paradis

## Abstract

**Objective:** The transcriptomic classification of intrahepatic cholangiocarcinomas (iCCA) has been recently refined from two to five classes, associated with pathological features, targetable genetic alterations and survival. Despite its prognostic and therapeutic value, the classification is not routinely used in the clinic because of technical limitations, including insufficient tissue material or the cost of molecular analyses. Here, we assessed a self-supervised learning (SSL) model for predicting iCCA transcriptomic classes on whole-slide digital histological images (WSIs)

**Design:** Transcriptomic classes defined from RNAseq data were available for all samples. The SSL method, called Giga-SSL, was used to train our model on a discovery set of 766 biopsy slides (n=137 cases) and surgical samples (n=109 cases) from 246 patients in a five-fold cross-validation scheme. The model was validated in The Cancer Genome Atlas (TCGA) (n= 29) and a French external validation set (n=32).

**Results:** Our model showed good to very good performance in predicting the four most frequent transcriptomic class in the discovery set (area under the curve [AUC]: 0.63-0.84), especially for the hepatic stem-like class (37% of cases, AUC 0.84). The model performed equally well in predicting these four transcriptomic classes in the two validation sets, with AUCs ranging from 0.76 to 0.80 in the TCGA set and 0.62 to 0.92 in the French external set.

**Conclusion:** We developed and validated an SSL-based model for predicting iCCA transcriptomic classes on routine histological slides of biopsy and surgical samples, which may impact iCCA management by predicting prognosis and guiding the treatment strategy.

## INTRODUCTION

Intrahepatic cholangiocarcinoma (iCCA) is the second most common primary liver cancer; the incidence is increasing worldwide and the prognosis is poor [1,2]. Although surgery is the only curative treatment for iCCA, only 20-40% of patients can benefit from surgery because of diagnosis at an advanced stage [3,4]. For disease not able to be treated by surgery, locoregional treatment in non-metastatic intrahepatic cases or systemic treatment (gemcitabine and cisplatin +/-durvalumab or pembrolizumab in first line) in metastatic cases is proposed, with a median overall survival of 15 and 12 months, respectively [5–8].

Recent advances in the pathobiological and molecular understanding of iCCA have provided prognostic and theranostic factors for a better clinical management of iCCA. From a transcriptomic point of view, for a long time, two distinct groups of iCCA have been identified: an inflammatory class (40% of cases) characterized by activation of inflammatory signalling pathways, and a proliferation class (60% of cases) characterized by the activation of oncogenic signalling pathways [9]. Recently, this classification has been refined into five classes. The inflammatory class has been divided into two sub-classes (inflammatory stroma and immune classical) and the proliferative class into three subclasses, namely hepatic stem-like, tumour classical and desert-like. Of note, this classification has been associated with tumour microenvironment composition, genetic alterations and prognosis and could guide the treatment strategy [10]. In particular, the hepatic stem-like class, the most frequent transcriptomic class, has been associated with a better prognosis and targetable genetic alterations including *isocitrate dehydrogenase 1* (*IDH1)* mutations (observed in 16% of hepatic stem-like cases compared to 7% in other classes) and *fibroblast growth factor receptor 2* (*FGFR2)* fusions (observed in 13% of hepatic stem-like cases compared to 5% in other classes), particularly clinically relevant, because specific inhibitors (ivosidenib and pemigatinib, respectively) have been approved by the US Food and Drug Administration as second-line treatments for locally advanced or metastatic iCCA [11–13]. Moreover, the two inflammatory classes may particularly benefit from immunotherapy. This comprehensive transcriptomic classification is not used in routine practice because it is currently based on sophisticated molecular biology techniques (expensive, only accessible in expert centres) and requires adequate tissue samples rich in tumour cells.

Artificial intelligence (AI) models, particularly deep neural networks are rapidly emerging in the medical field, especially in imaging [14–17]. With the development of digital pathology and wide access to digitised whole slide images (WSI), AI approaches can be used for classification tasks, as for example distinguishing cholangiocarcinoma from secondary forms of liver metastatic adenocarcinoma [18]. AI approaches can also be used to identify prognostic microscopic features and transcriptomic classification in hepatocellular carcinoma [19–23]. Despite these successes, deep learning techniques require large datasets (>1000 slides) [24] and heavy computational machinery that limits in-depth studies on stratified datasets. We recently introduced Giga-SSL, a self-supervised learning (SSL) algorithm designed to generate generalist low-dimensional feature vectors of WSIs, which offers both computational efficiency and label-efficiency [25]. Moreover, the significance of the slide types (coloration, slides directly or not associated with molecular analysis, etc) used for predicting molecular alterations and the potential heterogeneity they may introduce has been insufficiently explored [26].

The main objective of the present study was to predict iCCA transcriptomic classes on WSI using the Giga-SSL model, with a focus on identifying the hepatic stem-like class (the most frequent class).

## METHODS

### Patient and samples

The workflow of the study is summarised in figure 1. For the discovery set, we selected 246 formalin-fixed paraffin-embedded (FFPE) iCCA cases (109 surgical specimens and 137 biopsies) archived during 2000 and 2021 in the pathology department of Beaujon Hospital (Clichy, France). The material represented 769 hematein eosin saffron (HES) slides, divided into 5 folds at the patient level for cross-validation. All available slides for the surgical cases (including preoperative biopsies when available, n=25) were selected for the study (median WSI per case: 5 [range 1-12]). The slides were scanned at 20x magnification with an Aperio scanner (ScanScope AT Turbo).

**Figure 1.**
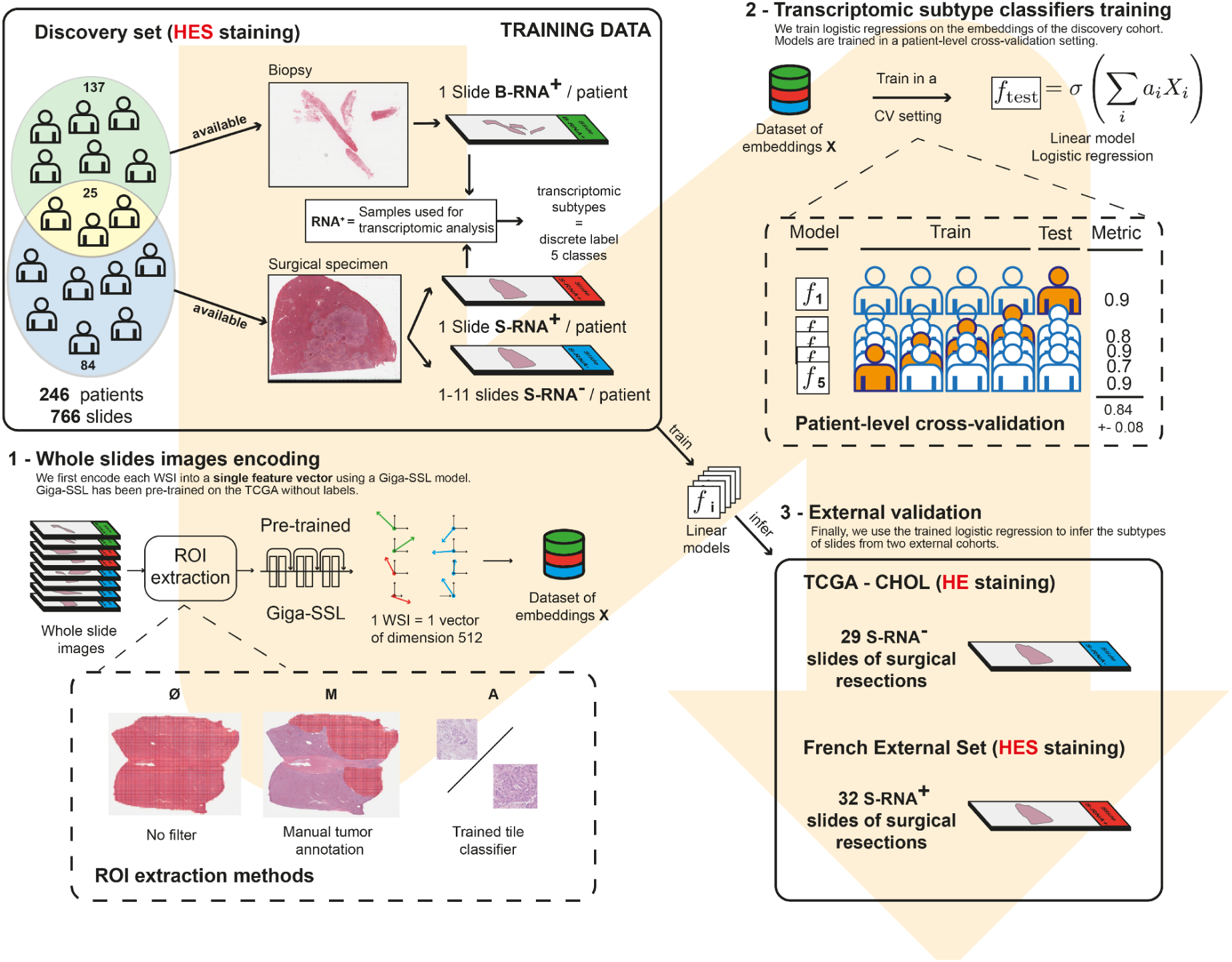
Flow-chart of the study. The model was first trained for predicting the five transcriptomic classes in a discovery set of FFPE iCCA biopsy and surgical samples (n=246 patients, Beaujon Hospital, Clichy, France) in a five-fold cross-validation scheme according to three different ROI extraction methods. It was then validated in a French external validation set (n=32 patients, Henri Mondor Hospital, Créteil, France) and a set of slides from TCGA (n=29 patients). *Formalin-fixed paraffin-embedded, FFPE; Hematein eosin, HE; Hematein eosin saffron, HES; cross-validation, CV; Region of interest, ROI; Slide from biopsy sample corresponding to the sample used for the transcriptomic analysis, slide B RNA+; Slide from surgical sample corresponding to the sample used for the transcriptomic analysis, slide S RNA+ ; Slide from surgical sample, not corresponding to the sample used for the transcriptomic analysis, slide S RNA-; Self-supervised learning, SSL; The cancer genome atlas, TCGA*.

For the external validation sets, we selected 29 iCCA cases (surgical FFPE samples, 29 WSI) from The Cancer Genome Atlas cholangiocarcinoma (TCGA-CHOL) public dataset and 32 iCCA surgical FFPE samples (32 WSI corresponding to the single most representative slide for each case) from the Pathology department of Henri Mondor Hospital (Créteil, France) (French external validation set).

We used the following selection criteria: 1) iCCA diagnosis after reviewing by an expert pathologist, 2) ≥1 available WSI from FFPE material, and 3) molecular analysis available.

Written consent was obtained from all patients as required by French legislation. This study was approved by the local ethics committee (IRB 00006477 CER Paris Nord no. CER-2022-168).

The clinical and biological data recorded were age at surgery, sex, risk factors of iCCA, tumour size (radiological assessment for biopsy and pathological assessment for surgical specimens), number of tumours and overall survival.

### Pathology review

All histological slides were reviewed by an expert liver pathologist (AB) and the assessed tumour features are listed in Table S1 (Figure S1). The stage of fibrosis in the non-tumoral liver when available was evaluated according to the METAVIR staging system [27].

### RNA sequencing

#### RNA extraction

RNA sequencing was performed on the FFPE block selected for surgical specimens, specifically the most representative slide in the discovery and French external sets. These slides directly associated with transcriptomic analysis (consecutive slides) are labelled as surgical slides (S RNA+) whereas slides from other blocks indirectly associated with transcriptomic analysis in the discovery set and the TCGA set are labelled (S RNA-). For biopsies, the FFPE block used for RNA sequencing corresponded directly to the slide selected (labelled slide B RNA+) (Figure 1, Figure S2).

Briefly, 5 µm-thick sections with macrodissection as needed were cut from FFPE blocks. Total RNAs was further isolated by using the Qiagen FFPE RNA extraction kit (RNeasy FFPE kit, Qiagen) for the discovery set and the Recover AllTM Total Nucleic Acid Isolation Kit for the French external validation cohort (Invitrogen, Thermo Fisher Scientific).

#### Gene expression analysis

Gene expression was analysed by using the SMARTer Stranded Total RNA-Seq Kit for the discovery set and QuantSeq 3’ mRNA-seq Kit for the French external validation set. Only genes quantified in at least 50% of samples were retained for the analysis. Gene expression profiles were quantile-normalized. The mean expression of each gene-set-defined gene signature was computed following a gene-wise centring in each dataset (without variance scaling). The transcriptomic class with the highest gene-set averaged expression was assigned to each sample. The same process was used with the TCGA dataset.

### Slide preprocessing and tessellation

Slides from the discovery set were stained with HES and encoded in svs format. Slides from the external French validation set were stained with HES and encoded in ndpi format. Slides from the TCGA validation set were stained with hematoxylin-eosin (HE) and encoded in svs format. Tissue regions automatically extracted using Otsu thresholding were then exhaustively split into 2899811 patches of 224×224 pixels (without overlapping) at 10x using the OpenSlide library in Python.

We present the results in the discovery set according to the following three pre-processing protocols with or without extraction of the region of interest (ROI), each requiring varying levels of expert pathologist involvement (Figure 1, Figure S3):

1. No-filter (Ø): all tiles including tumour and non-tumour are processed as they are (encompassing both tumour and non-tumour regions).
2. Manual-filter (M): an expert pathologist (AB) extensively annotates tumour regions using ImageScope software, from which patches are extracted.
3. Learning-filter (A): tiles are filtered using logistic regression trained on a dataset of 3000 tile embeddings, randomly extracted and labelled by an expert pathologist (AB).

### Machine learning algorithms

#### Data split

Training involved using a 5-fold cross-validation framework in the discovery set. Splits were stratified according to the output variable, at the patient level. They were shared among all trainings to ensure the fairness of comparison.

#### Giga-SSL representations

The Giga-SSL model was trained on a single V100 GPU on the TCGA-FFPE dataset following the training framework provided in our previous study [25] with the exception of the following:

- Training was performed for 100 hours, or 7800 epochs
- WSI embeddings were ensembled over 100 views, then L2-normalized.

Finally, we used L2-regularised logistic regressions (C=7, max_iter = 10000, and class_weight set as “balanced”) as end classification models.

#### MIL baseline algorithms

Beside the Giga-SSL based classifications, we provided some baseline classification algorithms for comparison. They were based on the deep attention multiple instance learning (MIL) algorithm introduced in the work of Ilse *et al*[28] and slightly modified in Lazard *et al* [29]. MIL models were trained from scratch and operate on tile embeddings extracted from the last layer of pretrained ResNet18 [30]. We used ResNet18 pre-trained on imagenet and on the TCGA with MoCo (i.e the tile-encoder used in the Giga-SSL model).

#### External dataset inference

The probabilities predicted by the five trained logistic regression on the training set (each corresponding to a training fold) were averaged and performances were computed using these pooled probabilities.

### Statistical analysis

Continuous variables were compared by the use of Student’s t -test, and categorical variables were compared by use of chi-squared or Fisher’’s exact tests. Survival curves were estimated by the Kaplan-Meier method, compared with log-rank statistics. Pp <0.05 was considered statistically significant (SPSS software). The performance of AI models was assessed thanks with the area under the receiver operating characteristic curve (AUC) score, balanced accuracy score and F1 score (macro-average).

## RESULTS

### Patient characteristics

The main clinical and pathological features of patients and tumours for each dataset are in Table 1. The three datasets were similar for most clinical and pathological features, particularly age (mean 63 years [discovery and French external validation sets] and 64 years [TCGA set], p=0.810) and sex distribution (male, 57%, 72% and 45%, p=0.99). In the discovery and French external validation sets, the main risk factors were chronic alcohol intake (18% and 19%, p=0.859) and metabolic syndrome (30% and 16%, p=0.141). At the pathology level, the French external validation set had higher proportions of large duct tumours and intense immune tumour infiltration than the two other sets (22% *vs* 7% and 3%, p=0.018; 59% *vs* 38% and 33%, p=0.011).

**Table 1.**
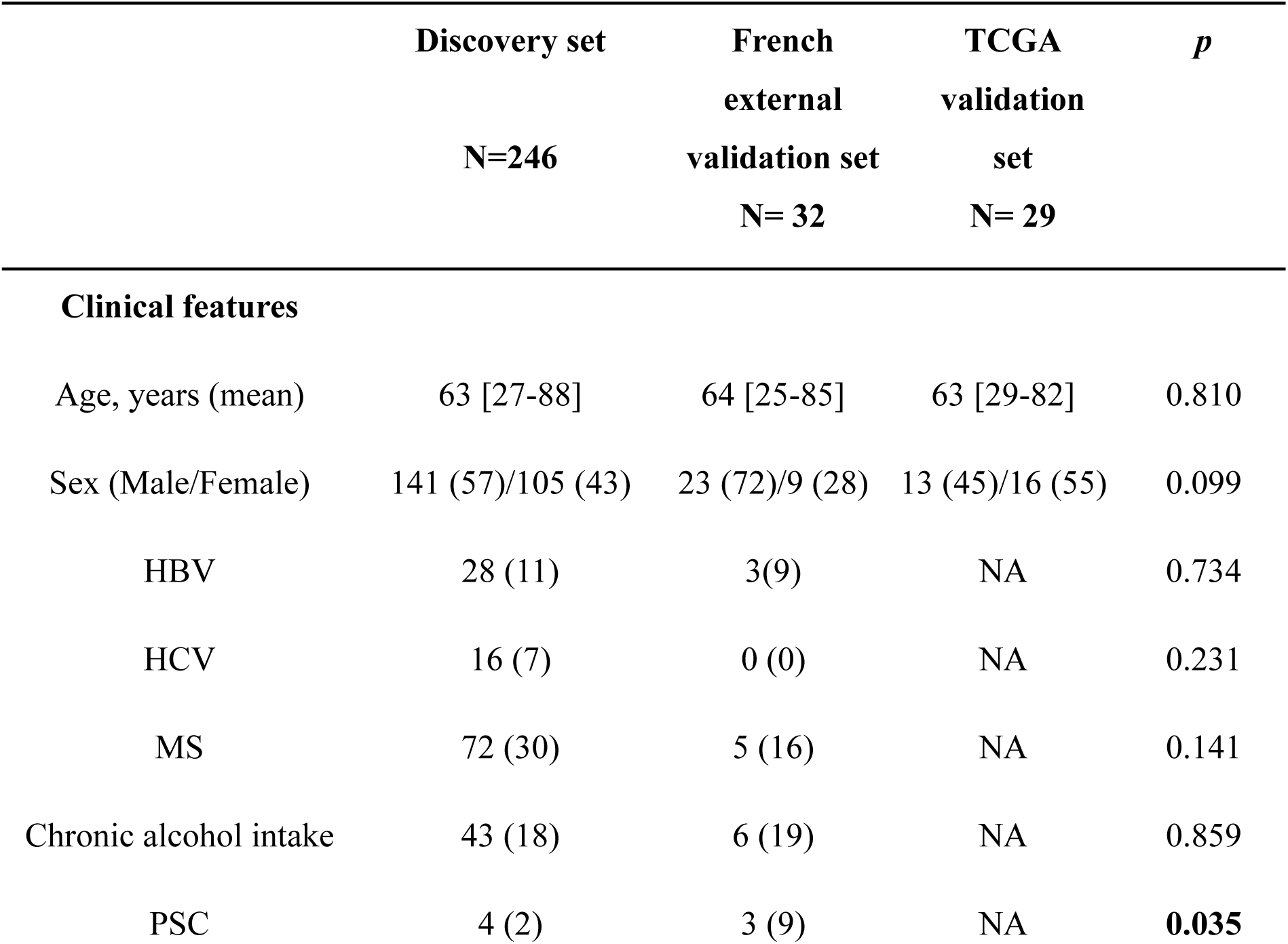
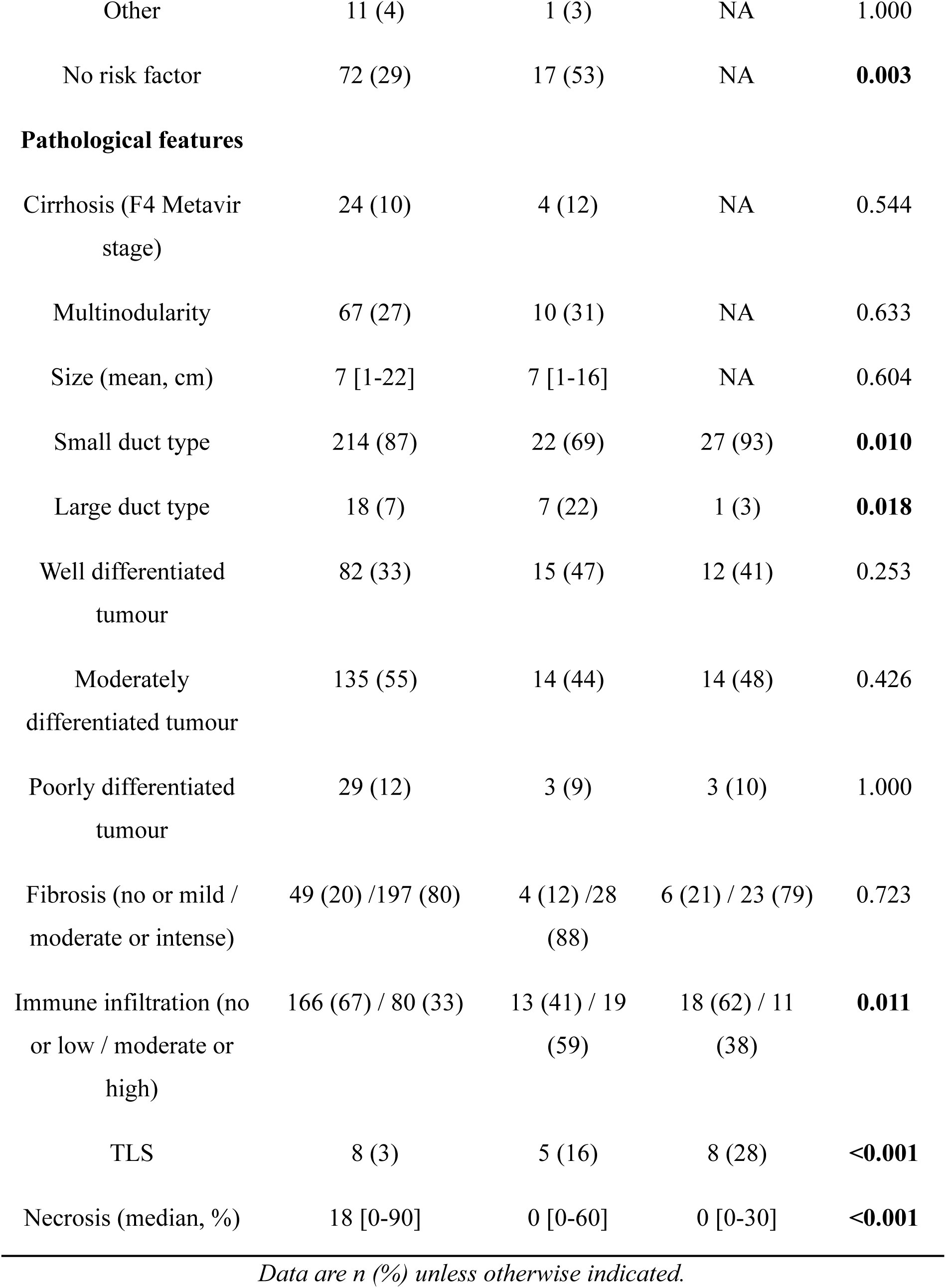
Clinical and pathological features of the different datasets of the study. Data not available, NA; hepatitis virus B, HBV; Hepatitis virus C, HCV; Metabolic syndrome, MS; Primary sclerosing cholangitis, PSC; Tertiary lymphoid structures, TLS. In caseIf of not available in the TCGA set, the statistical analyses were performed only between only the discovery and the French external validation sets.

|  | Discovery set<br>N=246 | French<br>external<br>validation set<br>N= 32 | TCGA<br>validation<br>set<br>N= 29 | *p* |
| --- | --- | --- | --- | --- |
| **Clinical features** |  |  |  |  |
| Age, years (mean) | 63 [27-88] | 64 [25-85] | 63 [29-82] | 0.810 |
| Sex (Male/Female) | 141 (57)/105 (43) | 23 (72)/9 (28) | 13 (45)/16 (55) | 0.099 |
| HBV | 28 (11) | 3(9) | NA | 0.734 |
| HCV | 16 (7) | 0 (0) | NA | 0.231 |
| MS | 72 (30) | 5 (16) | NA | 0.141 |
| Chronic alcohol intake | 43 (18) | 6 (19) | NA | 0.859 |
| PSC | 4 (2) | 3 (9) | NA | **0.035** |
| Other | 11 (4) | 1 (3) | NA | 1.000 |
| No risk factor | 72 (29) | 17 (53) | NA | **0.003** |
| **Pathological features** |  |  |  |  |
| Cirrhosis (F4 Metavir stage) | 24 (10) | 4 (12) | NA | 0.544 |
| Multinodularity | 67 (27) | 10 (31) | NA | 0.633 |
| Size (mean, cm) | 7 [1-22] | 7 [1-16] | NA | 0.604 |
| Small duct type | 214 (87) | 22 (69) | 27 (93) | **0.010** |
| Large duct type | 18 (7) | 7 (22) | 1 (3) | **0.018** |
| Well differentiated tumour | 82 (33) | 15 (47) | 12 (41) | 0.253 |
| Moderately differentiated tumour | 135 (55) | 14 (44) | 14 (48) | 0.426 |
| Poorly differentiated tumour | 29 (12) | 3 (9) | 3 (10) | 1.000 |
| Fibrosis (no or mild / moderate or intense) | 49 (20) / 197 (80) | 4 (12) / 28 (88) | 6 (21) / 23 (79) | 0.723 |
| Immune infiltration (no or low / moderate or high) | 166 (67) / 80 (33) | 13 (41) / 19 (59) | 18 (62) / 11 (38) | **0.011** |
| TLS | 8 (3) | 5 (16) | 8 (28) | **<0.001** |
| Necrosis (median, %) | 18 [0-90] | 0 [0-60] | 0 [0-30] | **<0.001** |

*Data are n (%) unless otherwise indicated.*

### Transcriptomic classes

The proportion of each transcriptomic class in each dataset is represented in Figure 2 and Table S2. The most frequent transcriptomic class was the hepatic stem-like class: 37% (90/246) of cases in the discovery set, 44% (14/32) in the French external validation set and 59% (17/29) in the TCGA validation set. The desert like class was very rare, observed in only 5% of cases (n=14). Of note, the distribution of the five transcriptomic classes differed between surgical and biopsy samples in the discovery set (Table S3). The hepatic stem-like class was more frequent in surgical than biopsy samples (49% *vs* 27%, p<0.001), whereas the immune classical class was more frequent in biopsy samples (31% vs 14%, p<0.002).

**Figure 2.**
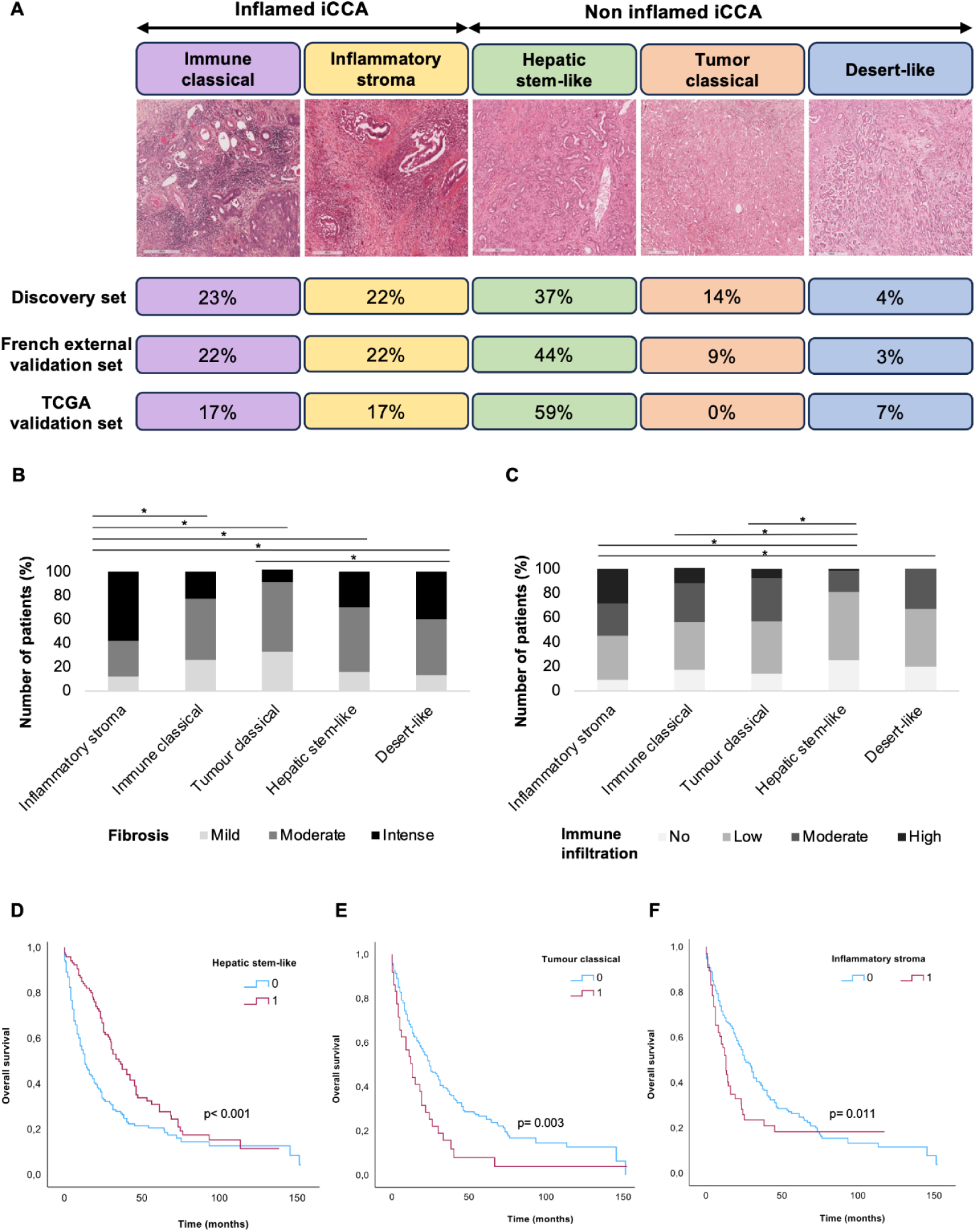
Distribution and characterisation according to histological features and overall survival of the five transcriptomic classes. (A) Distribution of each transcriptomic class according to the different datasets used in the study and representative histological images of each transcriptomic class (HES), (B) semi-quantitative assessment of the amount of tumour fibrosis in each transcriptomic class (*p<0.05), (C) Semi quantitative assessment of the amount of tumour immune infiltration in each transcriptomic class (*p<0.05), Kaplan-Meier curves for overall survivall according to (D) hepatic stem-like, (E) tumour classical and (F) inflammatory stroma transcriptomic class.

As expected, transcriptomic classes in all cohorts were associated with pathological features (Figure 2B-C). Marked tumour fibrosis was mainly observed in the inflammatory stroma class (n=38/66, 58%). A high tumour immune infiltration was predominant in inflammatory stroma and immune classical classes (n=19/66, 29% and n=8/69, 16%). Hepatic stem-like and desert-like classes exhibited low tumour immune infiltration (n=67/120, 56% and n=7/15, 47%). The five transcriptomic classes did not differ by tumour histological type (small duct *vs* large duct) (Figure S4).

At the clinical level, overall survival was significantly associated with three transcriptomic classes in all cohorts. It was improved with the hepatic stem-like class versus the other transcriptomic classes (overall survival median: 49 *vs* 35 months, hazard ratio [HR] 0.58; 95% confidence interval [CI] 0.44-0.75; p<0.001) but was altered with tumour classical and inflammatory stroma classes versus other classes (overall survival median: 21 *vs* 43 months, HR 1.76; 95% CI 1.09-2.82; p=0.003 and 31 *vs* 43 months, HR 1.50; 95% CI 1.04-2.17; p=0.011) (Figure 2D-F).aurel

### Using self-supervised WSI representations for transcriptomic class prediction

We initially focused on the binary classification task of the hepatic stem-like class, the most frequent class, before expanding our analysis to other transcriptomic classes, apart from the desert-like class, which was too rare in our three sets to develop a predictive model for this class.

### Predicting the hepatic stem-like class

#### In the discovery set

Table 2 shows cases of the cross-validated performances of both the Giga-SSL and MIL models in the discovery cohort for the hepatic stem-like binary classification task. The performance of the Giga-SSL model peaked when combined with manual tumour annotation, which resulted in a mean AUC of 0.84.

**Table 2.** Cross-validated performance of the Giga-SSL and multiple instance learning (MIL) models on the discovery cohort for the hepatic stem-like binary classification task according to three different pre-processing protocols. Protocols: learning filter (A): tiles were are filtered using logistic regression trained on a dataset of 3000 tile embeddings, randomly extracted and labelled by an expert pathologist; manual filter (M): an expert pathologist extensively annotates tumour regions using ImageScope software, from which patches are extracted; no filter (Ø): all tiles including tumour and non-tumour are processed as they are (encompassing both tumour and non-tumour regions). *Area under the curve, AUC; Learning-Filter, A; Manual-Filter, M; No-Filter,* Ø*; Multiple intance learning, MIL; Self-supervised learning, SSL*.

| Model | Tile Filter | AUC score | Balanced<br>accuracy score | F1 score |
| --- | --- | --- | --- | --- |
| **GigaSSL** | ∅ | 0.8 | 0.72 | 0.72 |
|  | M | **0.84** | **0.76** | **0.76** |
|  | A | 0.82 | 0.74 | 0.75 |
| **MoCo + MIL** | ∅ | 0.74 | 0.67 | 0.67 |
|  | M | 0.82 | 0.75 | 0.75 |
|  | A | **0.82** | **0.75** | **0.74** |

Indeed, performance improved when a protocol of ROI extraction was applied to WSIs, regardless of whether the extraction of the tumour tiles was manual or learned. This improvement was particularly notable when using the classic MIL models, with the absence of WSI filtering leading to an 8-point decrease in AUC. For the Giga-SSL models, the absence of WSI filtering led to a 4-point decrease in AUC.

Among surgical cases with several slides (n=106, 626 slides), the mean variance in prediction per case was 0.147 (range 0.007-0.418). In 51% of cases (54/108), the prediction was the same on each of the slides for the case.

#### External validation of the model for predicting the Hepatic stem-like class

Table 3 presents the results of the external validation of models trained on all slides of the discovery cohort, with a manual filter applied to the Whole Slide Images (WSIs). Logistic regression trained on the Giga-SSL embeddings of the discovery cohort demonstrated strong transferability to both the French external set (AUC=0.86) and TCGA sets (AUC=0.76). Notably, the TCGA cohort slides were stained with HE, which differs from the staining protocol used for the discovery cohort’s slides (HES) and further emphasizes the generalizability of the models.

**Table 3.** Model performance on external validation sets. The evaluated model is the ensemble of the five models trained in the cross-validation setting on the discovery cohort. AUC scores AUC scores are averaged over folds as described in the Methods section. *Area under the curve, AUC; The cancer genome atlas, TCGA*.

| Validation dataset | AUC score | Balanced accuracy score | F1 score |
| --- | --- | --- | --- |
| TCGA set | 0.76 | 0.73 | 0.74 |
| French external set | 0.86 | 0.73 | 0.71 |

#### Influence of the pre-processing ROI extraction protocol for predicting the hepatic stem-like class

We detail in Supplementary Table 3 the external validation results when models were trained according to the three different pre-processing protocols with or without ROI extraction. As observed in the cross-validated experiments on the discovery set, using an ROI extraction method (extraction only of the tumour tiles) was advantageous for both external datasets versus extraction of both tumour and non-tumour tiles.

#### Effect of slide composition of the training set for predicting of the hepatic stem-like class

In Figure 1 and Figure S2, the discovery dataset is illustrated to contain various types of slides, directly (RNA+) or indirectly associated to transcriptomic analysis (RNA-), including both surgical slides (S) and biopsy (B) slides. We aimed to determine how the composition of the training set affected the generalisation performance of classification models. For this, we trained models using training sets with different compositions of S RNA+ (or S+), S RNA-(or S-) and B RNA+ (or B) slides and monitored the prediction performance on the two external validation sets. The results are in Figure 3.

**Figure 3.**
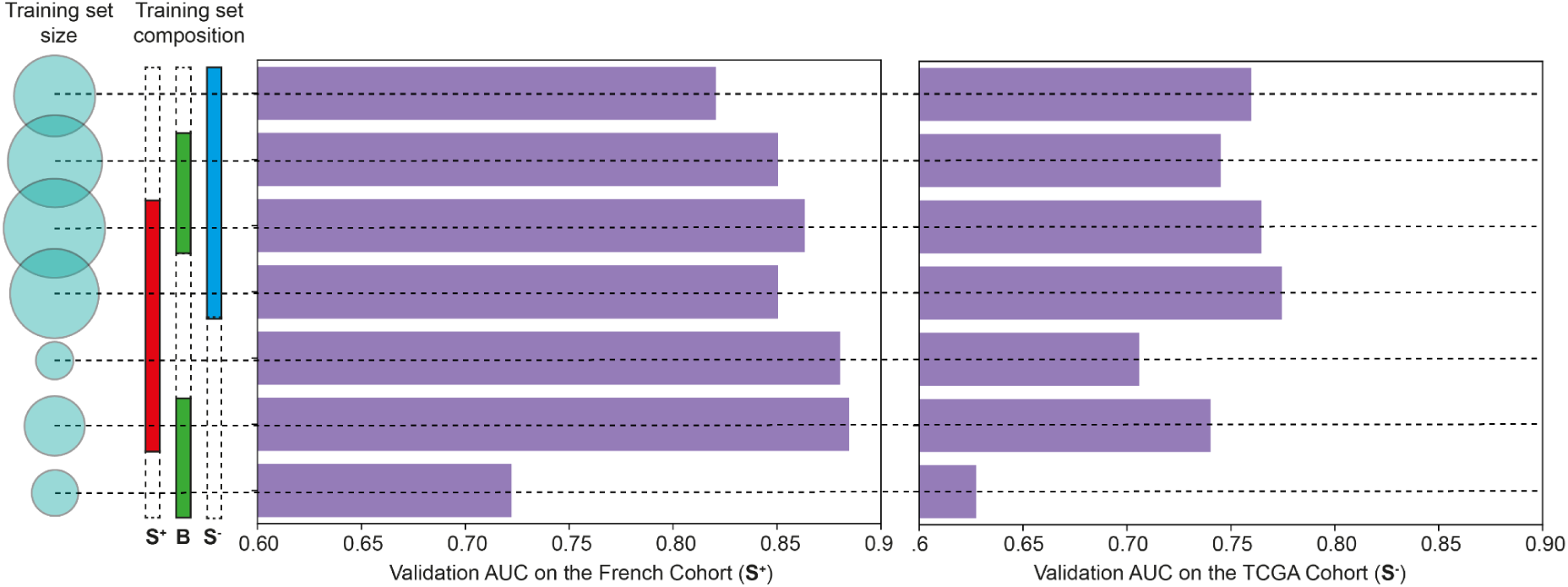
Effect of the composition of the training set on the performance of the Giga-SSL model for the hepatic stem-like binary classification task. Each row represents the performance of models trained on a specific subset of the discovery cohort. The characteristics of these subsets can be understood by looking at the intersection of the dotted line with the columns “training set size” and “training set composition”. For example, the third row represents models trained on the complete training dataset (S+, B, and S-), which indicates it as the largest training set. *Area under the curve, AUC; Self supervised learning, Slide from biopsy sample corresponding to the sample used for the transcriptomic analysis, B RNA+; Self-supervised learning, SSL; Slide from surgical sample corresponding to the sample used for the transcriptomic analysis,S RNA+; Slide from surgical sample, not corresponding to the sample used for the transcriptomic analysis, S RNA-; Region of interest, ROI*.

For the French external validation set, consisting solely of S RNA+ slides, incorporating S RNA-slides into the training set seemed detrimental to performance. Remarkably, the highest performance was achieved when the training set was limited to slides that were directly associated with the transcriptomic analysis (S RNA+ and B RNA+). Furthermore, performance was very close to that for the combined set (S RNA+ and B RNA+) when training solely with S RNA+ slides, despite being the smallest possible training dataset. Moreover, the addition of biopsy slides B RNA+ to surgical specimen slides slightly improved the validation performance.

Next, we turned to the validation on the TCGA dataset, exclusively consisting of S RNA-slides (i.e., slides for which the transcriptome was analysed on a different tissue block). For this dataset with a putatively noisier ground truth, the model’s performance seemed closely related to the size of the training data rather than its composition (Figure 3).

### Prediction of the other transcriptomic classes

We conducted analogous experiments for predicting other transcriptomic classes including inflammatory stroma, immune classical and tumour classical. As with the hepatic stem-like class, we trained binary classifiers for each class using a patient-level five-Fold cross-validation setting and then applied them to the validation sets. We manually set the ROI method and trained using the complete dataset (S RNA+, B RNA+, S RNA-). Table 4 presents the results of these experiments. The three classes could be predicted in a cross-validation setting and demonstrated generalisation capabilities on the validation sets.

**Table 4.** Predictions for the five transcriptomic classes. This table presents cross-validated (CV) and generalisation outcomes for both the French external and TCGA sets for the classification tasks related to the five transcriptomic classes. The TCGA cohort lacked any tumour classical samples. We trained the models using all slides (biopsies and surgical resections) and applied manual filter on the training and validation sets. Results are not presented when the metric is computed using less than two samples. *Cross-validation, CV; The cancer genome atlas, TCGA; Not applicable, NA*.

| Transcriptomic classes | CV results in the<br>discovery set | French external set<br>(N = 32) | TCGA set<br>(N = 29) |
| --- | --- | --- | --- |
| Hepatic stem-like | 0.84±0.06 | 0.86 (N=14) | 0.76 (N=17) |
| Tumour classical | 0.77±0.09 | 0.88 (N=3) | na (N=0) |
| Inflammatory stroma | 0.72±0.10 | 0.92 (N=7) | 0.80 (N=5) |
| Immune classical | 0.63±0.08 | 0.62 (N=7) | 0.78 (N=5) |

The classification task for the Inflammatory Stroma class outperforms the others on the validation set, achieving an AUC of 0.92 for the French set and 0.80 for the TCGA set. Despite the absence of tumour classical slides in the TCGA dataset, this class still seemed predictable and showed good generalisation with an AUC of 0.88 on the French set. However, despite some discernible signal for the Immune Classical class, it was the most challenging to classify.

## DISCUSSION

We show in this study that our SSL method applied to routine WSI is performant for predicting iCCA transcriptomic classes, particularly for hepatic stem-like and inflammatory stroma classes, associated with the prognosis and expected to impact on the treatment response to treatment, especially regarding immunotherapy and targeted therapies [9,10]. As previously described, the most frequent transcriptomic class in our three datasets was the hepatic stem-like class, which was associated with a better overall survival. We also confirmed in our cohorts the association between the two inflammatory classes and the tumour microenvironment composition (inflammation or/and fibrosis), which may benefit from immune checkpoint inhibitor treatments [10].

Our model showed good performance for predicting the transcriptomic classes particularly the hepatic stem-like, tumour classical and inflammatory stroma classes with AUCs of about 80% in the two external validation sets. The good predictions of these three transcriptomic classes are particularly interesting because they are all associated with overall survival, including in our cohorts and could impact the treatment strategy [10]. Furthermore, because the hepatic stem-like group is associated with targetable molecular alterations [10], our model could be used for pre-screening because even though recent guidelines recommend their investigation, molecular profiling may be challenging and cannot be carried out in routine practice, mainly for logistical and technical reasons [6].

Currently, the leading methods for WSI classification rely on MIL [28,31]. However, annotated datasets are often small, typically a few hundred to a few thousand WSIs, which may result in overfitting and underperforming models, and large unannotated datasets of tens of thousands WSI are available. Here, we used a slide level SSL model, called Giga-SSL, allowing us to leverage the large number of WSIs without annotations to infer powerful slide representations [25]. Our model surpassed the performance of the standard MIL model in the binary classification task for the hepatic stem-like class. We observed a gain of 2 to 6 points in AUC according to the pre-processing ROI extraction method. Besides a slight improvement in classification performance, this model significantly improves efficiency by increased speed and reduced use of computational resources. After the WSI embedding, all analyses used logistic regression and were seamlessly processed on a laptop CPU.

Moreover, our models could predict the transcriptomic classes on HES or HE slides, despite being exclusively trained on HES slides. This is particularly interesting because the routine staining performed differs among countries and laboratories leading to colour heterogeneity that could affect the performance of the AI model [32,33]. In addition, our model demonstrated better prediction when applied to tumour tiles rather than to all tiles (including non-tumour tiles) which suggests that the essential information regarding transcriptomic subclasses is contained in the tumour itself, rather than in its environment and other parts of the tissue. To bypass the time-intensive process of manual annotations by a pathologist, our data support the use of an automatic learning filter given its close performance to that when using a manual filter, which represents a favourable balance between time invested and classification performance.

Furthermore, we provide a comparative analysis that explores the influence of training set composition on prediction accuracy. Our findings suggest that intra-tumoral heterogeneity can negatively affect training when non-consecutive slides are used for molecular profiling and assessment of pathology. This discrepancy introduces label noise, in that the molecular class we aim to predict may not align with the tissue captured in the image. Despite a well-known requirement to have large datasets for effective neural network training, our results suggest that datasets with high-confidence labels outperform larger, noise-prone datasets.

Moreover, Kather *et al*[26] found that flash-frozen slides yielded better performance in molecular prediction tasks within the TCGA dataset, despite their poorer morphological quality compared with FFPE slides. This anomaly could also be attributed to label noise arising from tumour heterogeneity, because the molecular labels in TCGA are extracted from flash-frozen samples. These insights underscore the importance of using consecutive slides for molecular class prediction and could potentially inform the design of future studies. Finally, though more research is needed to validate the clinical use of such predictive models, patient stratification would benefit from combining the prediction of several WSIs sampled in different blocks, which would help capture the main iCCA class of the tumour and help in exploring the intratumour heterogeneity. Indeed, intratumour molecular heterogeneity in liver cancers, including iCCA, is a key feature that may explain treatment failure and patient prognosis[34]. Furthermore, our transcriptomic analysis of the three datasets revealed that certain cases exhibited a profile that spanned multiple transcriptomic classes, and classifying them into a single class is challenging.

Importantly, we included both biopsy and surgical samples in the discovery set to better reflect clinical practice. Even though using biopsies might have reduced our model’s performance during cross-validation, it improved performance on the French external validation set. Hence, biopsies may provide complementary information to surgical specimen WSIs (see Figure 3) and increase the robustness of the trained network. Currently, most AI studies of primary liver cancers have focused on surgical samples[16,19–21], but most patients with iCCA will not undergo surgery, which may introduce a selection bias. Of note, we found transcriptomic classes differentially represented between surgical and biopsy cases, which highlights the importance of working with both tissue specimen. Few studies, mainly focusing on diagnostic tasks, have laid the groundwork for using biopsies and have demonstrated that encouraging deep-learning–based results can be obtained in this type of sample despite their size[18,35,36]. Thus, our model could be a useful molecular screening tool particularly in the context of biopsy in which the molecular analysis is not always possible because of the low amount of materials.

Our study has some limitations. The proportion of each transcriptomic class was unbalanced and in particular the desert-like class was rare, representing less than 10% of cases in all sets, not allowing to develop a predictive model for this class. In addition, we were unable to evaluate the predictive performance of our model for the tumour classical class in the TCGA set, because of there were no cases of this class in this set. Learning from a larger number of cases could be beneficial, but finding complete datasets containing survival and transcriptomic data, and histological slides of FFPE iCCA samples is difficult, as evidenced by the low number of cases available in the TCGA dataset. Finally, the Giga-SSL models are not visually interpretable. However, the histological review performed beforehand and the results of Martin-Serrano *et al*[10] have allowed for highlighting different histological characteristics between transcriptomic classes, particularly in the tumour microenvironment composition.

## Conclusion

We have developed and validated an SSL model able to predict iCCA transcriptomic classes on routine WSIs from routine biopsy and surgical samples. This model showed good to very good performance for classifying hepatic stem-like, tumour classical, inflammatory stroma classes and immune classical classes. We have demonstrated the importance of the pre-processing method with ROI extraction and that training was more effective on a small dataset of slides directly associated with transcriptomic analysis (consecutive slides) than on a larger, noisier dataset (slides from other blocks indirectly associated with transcriptomic analysis). The ability to predict transcriptomic iCCA classes on routine WSIs could affect disease management by predicting prognosis and guiding the treatment strategy (immunotherapy in inflammatory classes or targeted molecular therapies in hepatic stem-like class).

### Abbreviations

AI: artificial intelligence
AUC: area under the curve
FFPE: formalin-fixed paraffin-embedded
FGFR2: fibroblast growth factor receptor 2
HE: hematoxylin-eosin
HES: hematein eosin saffron
iCCA: intrahepatic cholangiocarcinoma
IDH1: isocitrate dehydrogenase 1
MIL: multiple instance learning
SSL: self-supervised learning
TLS: tertiary lymphoid structures
TGCA: The Cancer Genome Atlas
TCGA-CHOL: The Cancer Genome Atlas-cholangiocarcinoma
WSI: whole slide image

## Acknowledgement

We acknowledge the iGenSeq core facility at ICM for the benefit of its equipment and services, and the Nuovo-Soldati foundation for its support.

## Supplementary material

**Table S1.** List of morphological criteria assessed by the expert pathologist for all cases of the three datasets.

| **Histological criteria** | **Assessment** |
| --- | --- |
| **Tumour grade** | Classification into well, moderately or poorly differentiated tumour according to the 5<sup>th</sup> edition of the WHO classification |
| **Tumour histological type** | Small duct, large duct or other subtypes |
| **Necrosis** | Percentage |
| **Tumour fibrosis** | Semi quantitatively assessed, classified into three classes: no or mild, moderate and intense |
| **Immune tumour infiltration** | Semi quantitatively assessed, classified into four classes: no inflammation, low, moderate and high |
| **Tertiary lymphoid structure (TLS)** | Presence or absence |

**Table S2.**
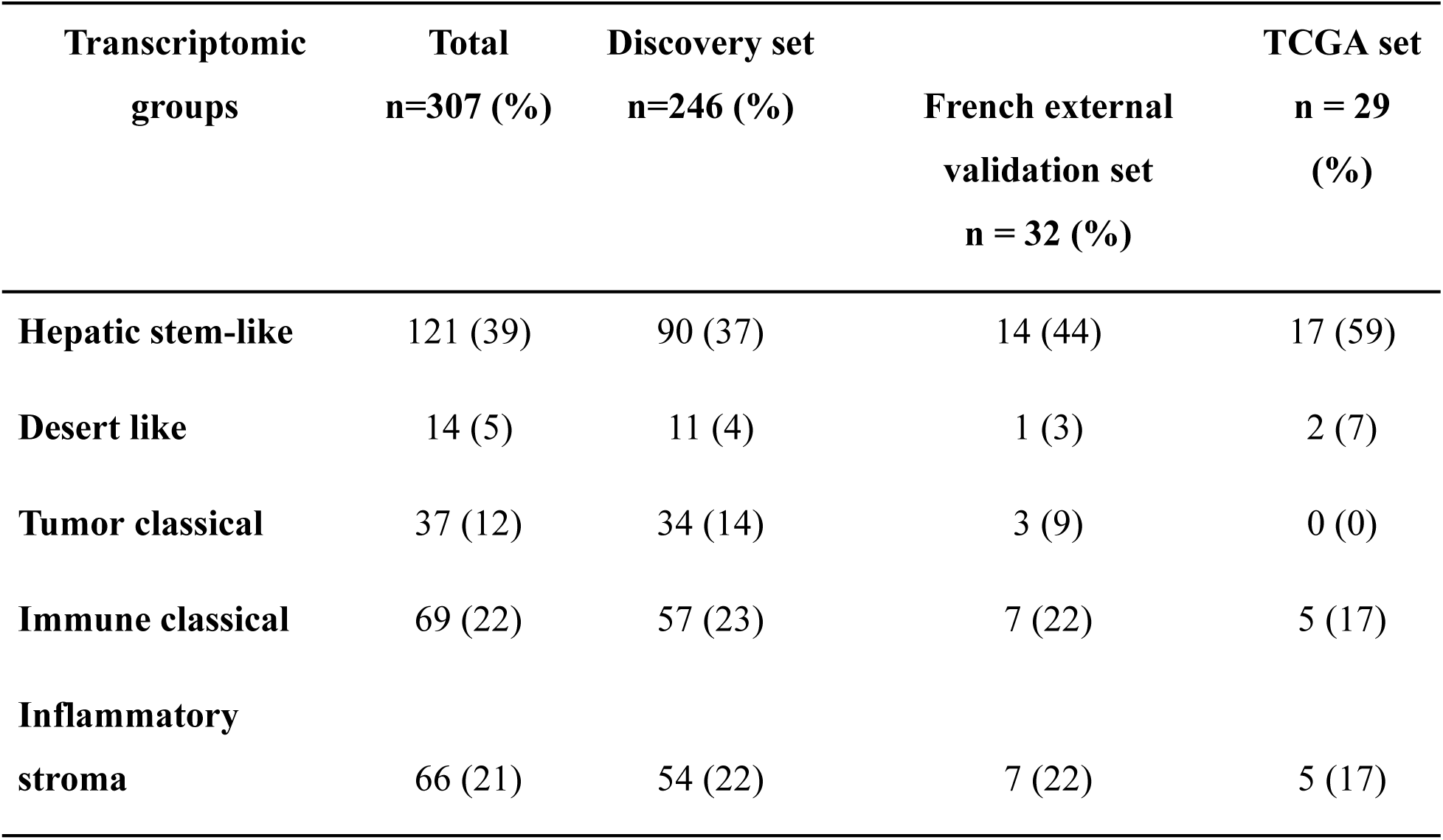
Repartition of the five transcriptomic classes in the discovery set and in the two external validation sets.

**Table S3.** Repartition of the five transcriptomic classes according to the type of samples.

| **Transcriptomic groups** | **Total**<br>**n=246 (%)** | **Surgical samples**<br>**n=109 (%)** | **Biopsies**<br>**n = 137 (%)** | ***p*** |
| --- | --- | --- | --- | --- |
| **Hepatic stem-like** | 90 (37) | 53 (49) | 37 (27) | **<0.001** |
| **Desert like** | 11 (4) | 4 (4) | 7 (5) | 0.759 |
| **Tumor classical** | 34 (14) | 16 (15) | 18 (13) | 0.853 |
| **Immune classical** | 57 (23) | 15 (14) | 42 (31) | **0.002** |
| **Inflammatory**<br>**stroma** | 54 (22) | 21 (19) | 33 (24) | 0.439 |

**Table S3.**
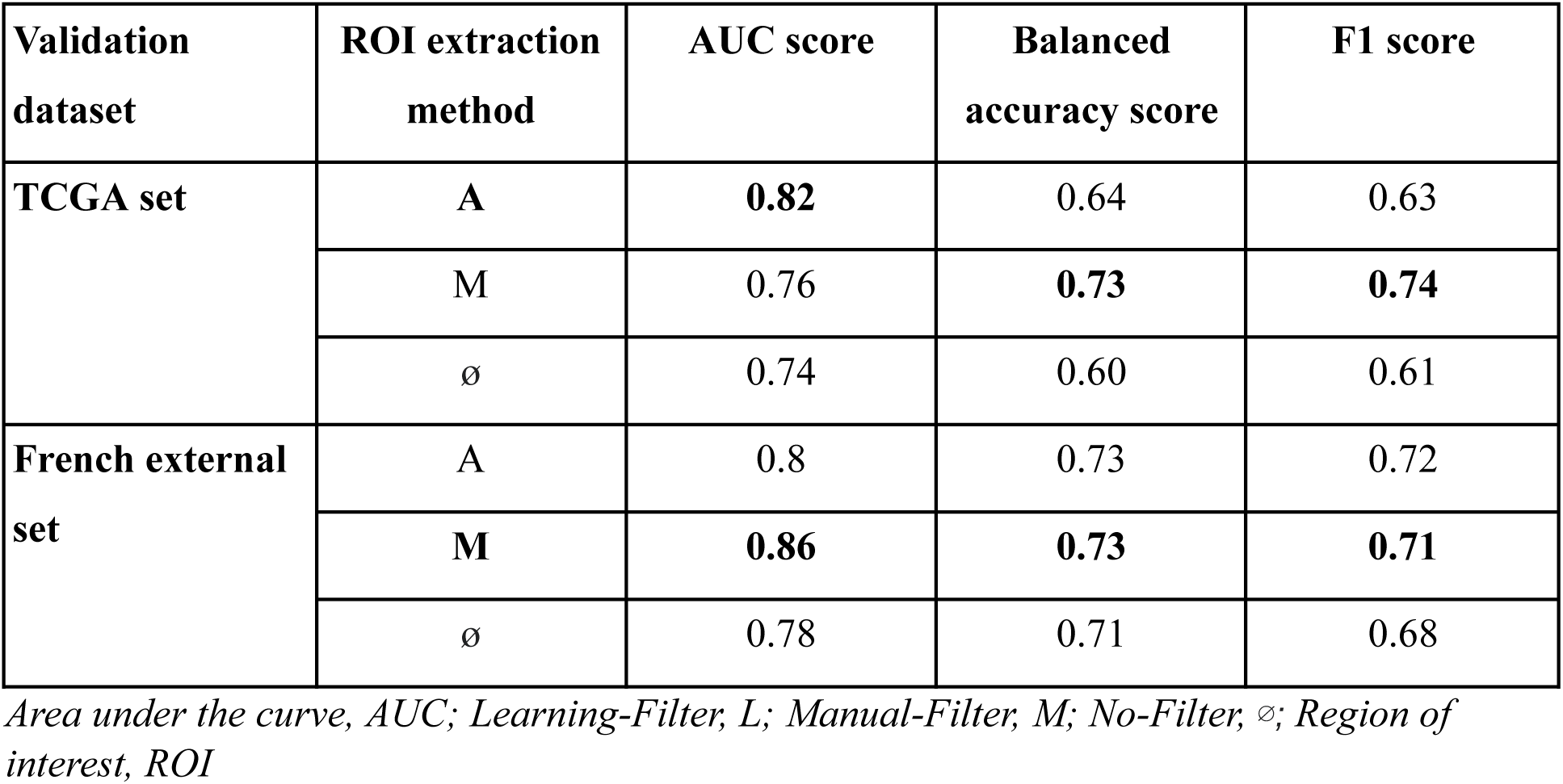
Effects of the ROI extraction method on the external validation performances for the Hepatic stem-like binary classification task. The same method is applied on both the training and validation datasets. On the TCGA set, no distinct advantage is observed for method A over M or vice versa, as it varies based on the metric under consideration. In the French external set, manually segmenting the tumour seems to be advantageous. Nevertheless, in both datasets, using an ROI extraction method is more effective than not using any.

| Validation dataset | ROI extraction method | AUC score | Balanced accuracy score | F1 score |
| --- | --- | --- | --- | --- |
| TCGA set | A | **0.82** | 0.64 | 0.63 |
|  | M | 0.76 | **0.73** | **0.74** |
|  | ∅ | 0.74 | 0.60 | 0.61 |
| French external set | A | 0.8 | 0.73 | 0.72 |
|  | **M** | **0.86** | **0.73** | **0.71** |
|  | ∅ | 0.78 | 0.71 | 0.68 |

*Area under the curve, AUC; Learning-Filter, L; Manual-Filter, M; No-Filter, ∅; Region of interest, ROI*

**Figure S1.**
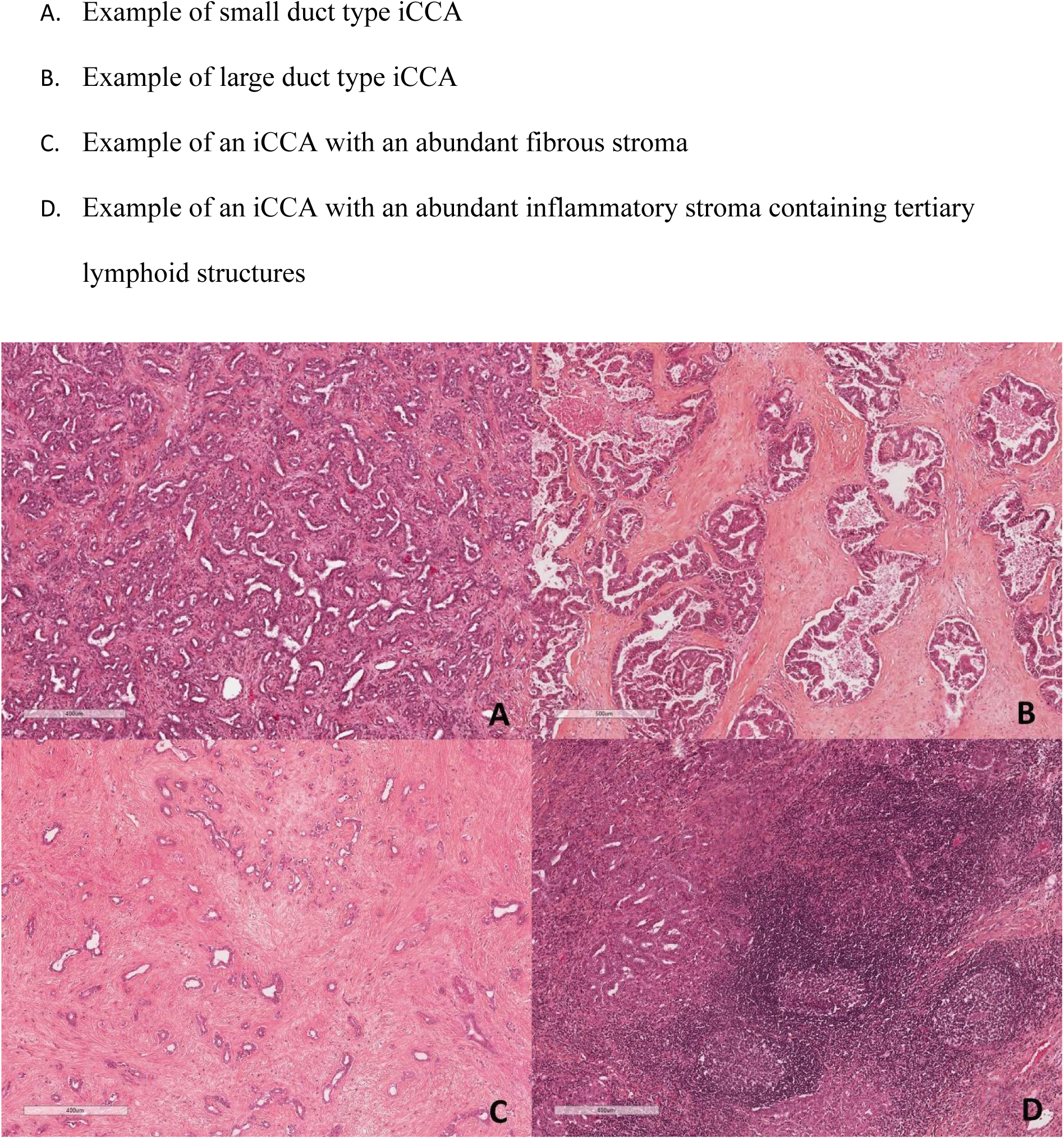
Histological features of iCCA.

**Figure S2.**
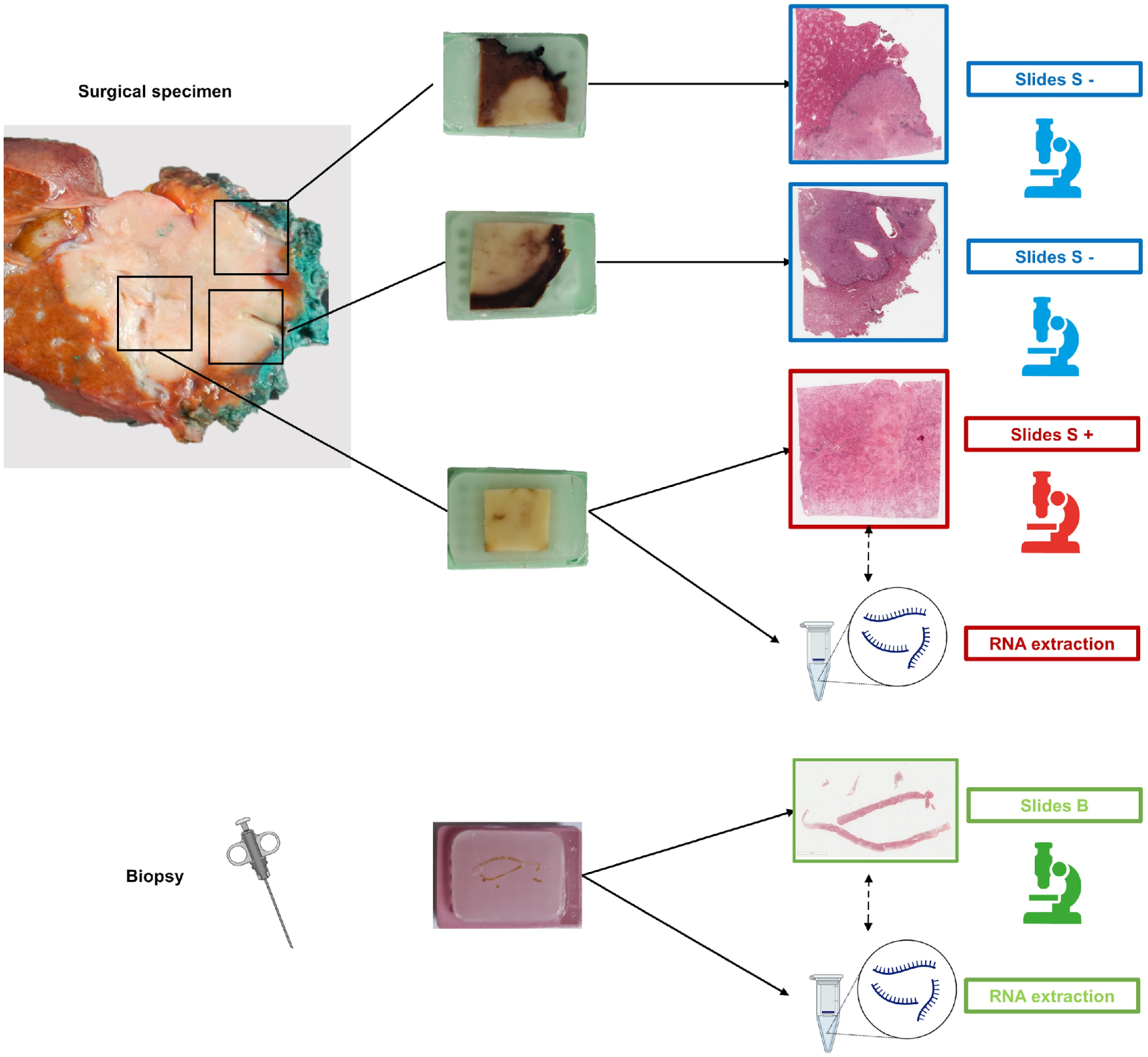
Type of slides including in the study. The slides directly associated with transcriptomic analysis (consecutive slides), have been labelled as surgical slides S+ whereas slides from other blocks indirectly associated with transcriptomic analysis in the discovery set and in the TCGA set have been labelled S-. For biopsy, the FFPE block used for RNA sequencing corresponded directly to the slide selected (labelled as slide B).

**Figure S3.**
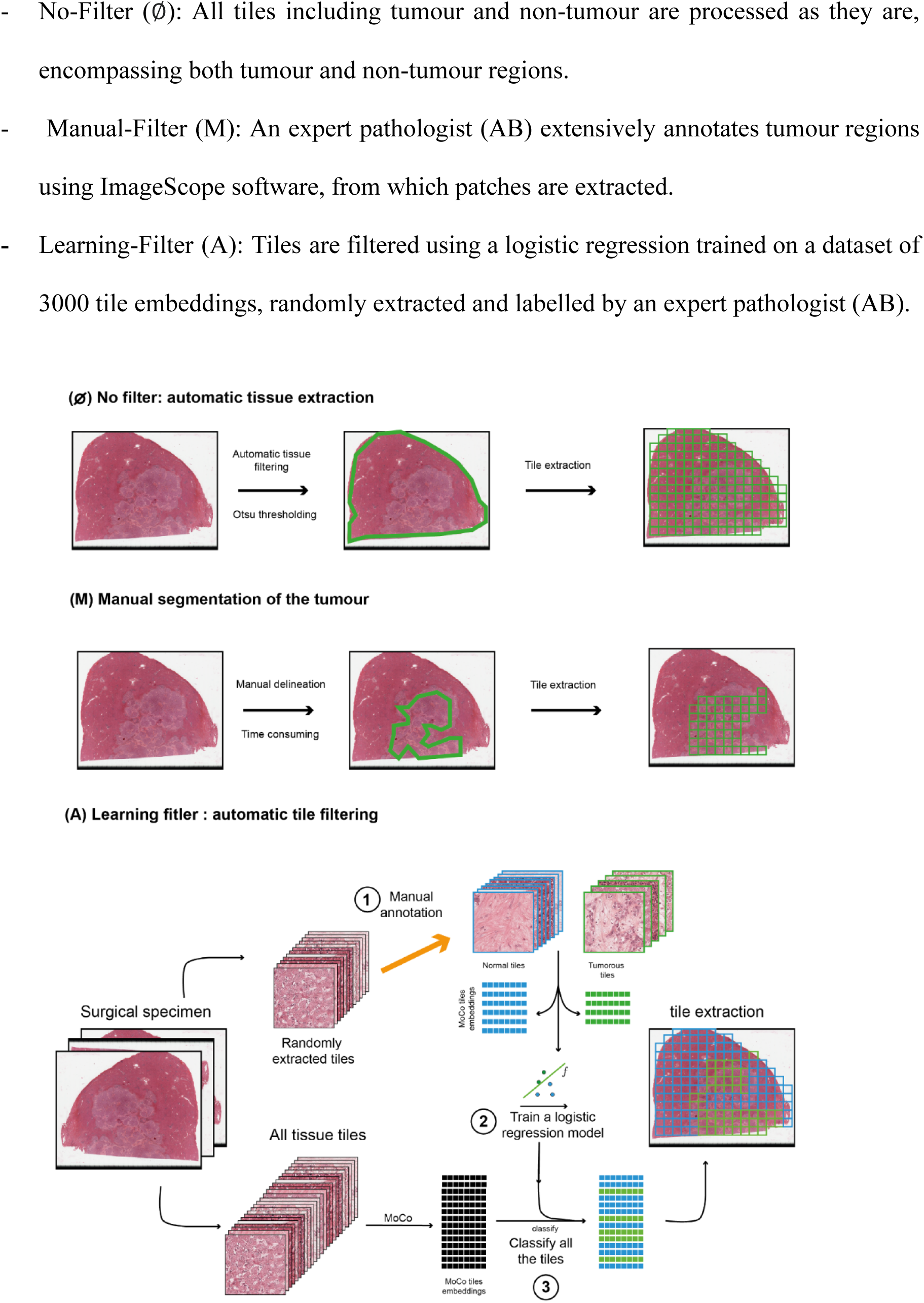
Three different pre-processing protocols with or without extraction of region of interest (ROI).

**Figure S4.**
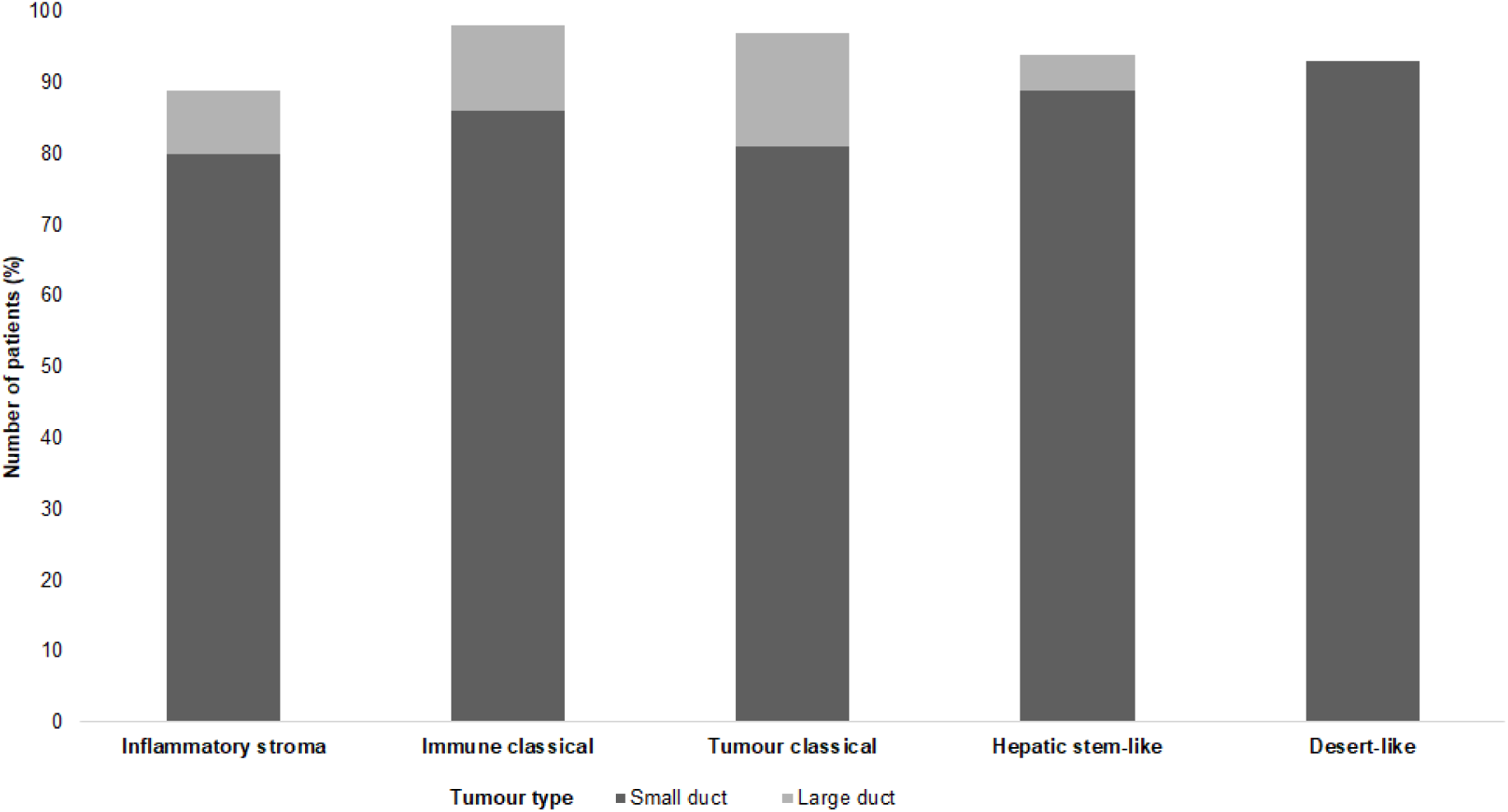
Histological iCCA subtypes (large vs small duct) according to the five transcriptomic groups.

## Notes

### Competing Interest Statement

The authors have declared no competing interest.

## REFERENCES

1 Global Burden of Disease Liver Cancer Collaboration, Akinyemiju T, Abera S, et al. The Burden of Primary Liver Cancer and Underlying Etiologies From 1990 to 2015 at the Global, Regional, and National Level: Results From the Global Burden of Disease Study 2015. JAMA Oncol. 2017;3:1683–91.

2 Rahnemai-Azar AA, Weisbrod A, Dillhoff M, et al. Intrahepatic cholangiocarcinoma: Molecular markers for diagnosis and prognosis. Surg Oncol. 2017;26:125–37.

3 Endo I, Gonen M, Yopp AC, et al. Intrahepatic cholangiocarcinoma: rising frequency, improved survival, and determinants of outcome after resection. Ann Surg. 2008;248:84–96.

4 Mavros MN, Economopoulos KP, Alexiou VG, et al. Treatment and Prognosis for Patients With Intrahepatic Cholangiocarcinoma: Systematic Review and Meta-analysis. JAMA Surg. 2014;149:565–74.

5 Bridgewater J, Galle PR, Khan SA, et al. Guidelines for the diagnosis and management of intrahepatic cholangiocarcinoma. J Hepatol. 2014;60:1268–89.

6 European Association for the Study of the Liver. Electronic address, European Association for the Study of the Liver. EASL-ILCA Clinical Practice Guidelines on Intrahepatic Cholangiocarcinoma. J Hepatol. 2023;S0168-8278(23)00185-X.

7 Oh D-Y, Ruth He A, Qin S, et al. Durvalumab plus Gemcitabine and Cisplatin in Advanced Biliary Tract Cancer. *NEJM Evid*. 2022;1:EVIDoa2200015.

8 Kelley RK, Ueno M, Yoo C, et al. Pembrolizumab in combination with gemcitabine and cisplatin compared with gemcitabine and cisplatin alone for patients with advanced biliary tract cancer (KEYNOTE-966): a randomised, double-blind, placebo-controlled, phase 3 trial. Lancet Lond Engl. 2023;401:1853–65.

9 Sia D, Hoshida Y, Villanueva A, et al. Integrative molecular analysis of intrahepatic cholangiocarcinoma reveals 2 classes that have different outcomes. Gastroenterology. 2013;144:829–40.

10 Martin-Serrano MA, Kepecs B, Torres-Martin M, et al. Novel microenvironment-based classification of intrahepatic cholangiocarcinoma with therapeutic implications. Gut. 2022;gutjnl-2021-326514.

11 Moeini A, Sia D, Bardeesy N, et al. Molecular Pathogenesis and Targeted Therapies for Intrahepatic Cholangiocarcinoma. Clin Cancer Res Off J Am Assoc Cancer Res. 2016;22:291–300.

12 Abou-Alfa GK, Macarulla T, Javle MM, et al. Ivosidenib in IDH1-mutant, chemotherapy-refractory cholangiocarcinoma (ClarIDHy): a multicentre, randomised, double-blind, placebo-controlled, phase 3 study. Lancet Oncol. 2020;21:796–807.

13 Abou-Alfa GK, Sahai V, Hollebecque A, et al. Pemigatinib for previously treated, locally advanced or metastatic cholangiocarcinoma: a multicentre, open-label, phase 2 study. Lancet Oncol. 2020;21:671–84.

14 Calderaro J, Seraphin TP, Luedde T, et al. Artificial intelligence for the prevention and clinical management of hepatocellular carcinoma. J Hepatol. 2022;76:1348–61.

15 Calderaro J, Kather JN. Artificial intelligence-based pathology for gastrointestinal and hepatobiliary cancers. Gut. 2021;70:1183–93.

16 Kather JN, Calderaro J. Development of AI-based pathology biomarkers in gastrointestinal and liver cancer. Nat Rev Gastroenterol Hepatol. 2020;17:591–2.

17 Chen X, Wang X, Zhang K, et al. Recent advances and clinical applications of deep learning in medical image analysis. Med Image Anal. 2022;79:102444.

18 Albrecht T, Rossberg A, Albrecht JD, et al. Deep learning-enabled diagnosis of liver adenocarcinoma. Gastroenterology. 2023;S0016–5085(23)04883-7.

19 Saillard C, Schmauch B, Laifa O, et al. Predicting survival after hepatocellular carcinoma resection using deep-learning on histological slides. Hepatol Baltim Md. Published Online First: 28 February 2020. doi: 10.1002/hep.31207

20 Zeng Q, Klein C, Caruso S, et al. Artificial intelligence predicts immune and inflammatory gene signatures directly from hepatocellular carcinoma histology. J Hepatol. 2022;77:116–27.

21 Cheng N, Ren Y, Zhou J, et al. Deep Learning-Based Classification of Hepatocellular Nodular Lesions on Whole-Slide Histopathologic Images. Gastroenterology. 2022;162:1948–1961.e7.

22 Shi J-Y, Wang X, Ding G-Y, et al. Exploring prognostic indicators in the pathological images of hepatocellular carcinoma based on deep learning. Gut. 2021;70:951–61.

23 Yamashita R, Long J, Saleem A, et al. Deep learning predicts postsurgical recurrence of hepatocellular carcinoma from digital histopathologic images. Sci Rep. 2021;11:2047.

24 Campanella G, Hanna MG, Geneslaw L, et al. Clinical-grade computational pathology using weakly supervised deep learning on whole slide images. Nat Med. 2019;25:1301–9.

25 Lazard T, Lerousseau M, Decencière E, et al. Giga-SSL: Self-Supervised Learning for Gigapixel Images. 2022. 10.48550/arXiv.2212.03273

26 Kather JN, Heij LR, Grabsch HI, et al. Pan-cancer image-based detection of clinically actionable genetic alterations. Nat Cancer. 2020;1:789–99.

27 Bedossa P, Poynard T. An algorithm for the grading of activity in chronic hepatitis C. The METAVIR Cooperative Study Group. Hepatol Baltim Md. 1996;24:289–93.

28 Ilse M, Tomczak JM, Welling M. Attention-based Deep Multiple Instance Learning.

29 Lazard T, Bataillon G, Naylor P, et al. Deep learning identifies morphological patterns of homologous recombination deficiency in luminal breast cancers from whole slide images. Cell Rep Med. 2022;3:100872.

30 He K, Zhang X, Ren S, et al. Deep Residual Learning for Image Recognition. 2015. 10.48550/arXiv.1512.03385

31 Li B, Eliceiri KW. Dual-stream Maximum Self-attention Multi-instance Learning. 2020. 10.48550/arXiv.2006.05538

32 Tellez D, Litjens G, Bándi P, et al. Quantifying the effects of data augmentation and stain color normalization in convolutional neural networks for computational pathology. Med Image Anal. 2019;58:101544.

33 Marini N, Otalora S, Wodzinski M, et al. Data-driven color augmentation for H&E stained images in computational pathology. J Pathol Inform. 2023;14:100183.

34 Ma L, Wang L, Khatib SA, et al. Single-cell atlas of tumor cell evolution in response to therapy in hepatocellular carcinoma and intrahepatic cholangiocarcinoma. J Hepatol. 2021;75:1397–408.

35 Pantanowitz L, Quiroga-Garza GM, Bien L, et al. An artificial intelligence algorithm for prostate cancer diagnosis in whole slide images of core needle biopsies: a blinded clinical validation and deployment study. Lancet Digit Health. 2020;2:e407–16.

36 Xu F, Zhu C, Tang W, et al. Predicting Axillary Lymph Node Metastasis in Early Breast Cancer Using Deep Learning on Primary Tumor Biopsy Slides. Front Oncol. 2021;11:759007.

